# Rational Design of Programmable Monodisperse Semi-Synthetic Protein Nanomaterials Containing Engineered Disulfide Functionality

**DOI:** 10.1101/2021.02.05.430015

**Authors:** Pavankumar Janardhan Bhandari, Britto S Sandanaraj

## Abstract

The reversible nature of disulfide functionality has been exploited to design intelligent materials such as nanocapsules, micelles, vesicles, inorganic nanoparticles, peptide and nucleic acid nanodevices. However, to our knowledge, there are no reports on design of semi-synthetic protein assemblies containing an engineered disulfide bond. Towards that end, herein, we disclose a new chemical methodology for the construction redox-sensitive protein assemblies using monodisperse facially amphiphilic protein-dendron bioconjugates. The disulfide functionality is strategically placed between the dendron and protein domains. The custom designed bioconjugates self-assembled into nanoscopic objects of a defined size dictated by the nature of dendron domain. The stimuli-responsive behavior of the protein assemblies is demonstrated using a suitable redox trigger.

## Introduction

Unlike standard covalent bonds, the disulfide bond is one of the most interesting covalent bonds. Because of this reason, developing chemical methodology to strategically place disulfide bond in both small molecules and macromolecules have gained enormous interest in the recent years.^1^ One of the most important applications of disulfide containing synthetic molecules or macromolecules is in the field of targeted drug delivery.^2^ The highly reducing environment of cytoplasm of a living cell (1-10 mM glutathione (GSH)) compared to the extracellular medium (20-40 μM) is effectively used to design smart drug delivery systems.^3^ To date, there are number of research groups working in this area to develop to disulfide containing amphiphilic polymers^4^, inorganic nanoparticles^5^, dendrimers^6^, virus-like particles^7^, peptide nucleic acids^8^ and nucleic acid nanostructures^9^. However, the engineering of disulfide bond in these systems is relatively easy as they are completely synthetic, soluble in organic solvent and therefore amenable for structural modification.

Globular proteins are emerging as nanoscale building blocks for the construction of smart drug delivery systems.^10^ Compared to other synthetic nanoscale building blocks, globular proteins are highly complex yet structurally more defined. Because of these reasons, there is a tremendous interest in the recent years for de novo design of stimuli-sensitive protein assemblies which are either completely synthetic or semi-synthetic.^11^ Engineering of native globular proteins through a chemical method yields semi-synthetic proteins with improved functions.^12^ However, most of the methods often yield polydisperse sample that are difficult to characterize and study. This is in sharp contrast to design of completely synthetic proteins which are made through genetic engineering/computational design.^13^ Both the approaches have proven to be powerful and have certain limitations and therefore complement each other. Our group initiated a research program in the area of de novo design of well defined semi-synthetic self-assembling proteins.^14^ To achieve this goal, we invented a new method named “Micelle-Assisted Protein Labeling (MAPLab) Technology”. This method provides opportunities to site-specifically install a synthetic hydrophobic group on to a native protein in a predictable manner.^14a^ We have demonstrated that this methodology can be used for labeling of large number of proteins with various synthetic hydrophobic small molecules^14a^, hydrophobic dendrimers^14b^ and monodisperse hydrophobic synthetic peptide^14c^. We have also demonstrated the design of de novo semi-synthetic proteins containing photo-sensitive moiety and studied their stimuli-responsive behavior in response to UV-light.^14a,d^ Athough, the use of a photosensitive group as an external trigger provides excellent spatiotemporal control. However, the phototoxicity associated with the UV light and its less penetration depth into the skin are the significant limitations.. On the other hand, use of internal trigger such as pH, redox and enzyme have been quiet successful for targeted delivery of drugs in humans.^15^

Strategic placement of disulfide functionality in a well defined semi-synthetic self-assembling protein is a challenging task for following reasons; (i) The quest for well defined system impose certain restriction on the use of macromolecular scaffolds, use of hydrophobic linear polymer is not an option because most of the synthetic polymers are polydisperse. Therefore, use of dendrimer is the only option as it can be synthesized as a single chemical entity with polydispersity index of 1.00. However the challenge is synthesis of a dendrimer is usually multi-step. (ii) secondly, hydrophobic dendrimer cannot be directly coupled to protein directly because of solubility issue and therefore hydrophilic monodisperse linker of appropriate length should be attached to the core of the dendrimer, again synthesis of monodisperse linker increases of number of synthetic steps overall. (iii) Finally, dendrimer containing monodisperse linker should be solubilized in an aqueous medium for site-specific bioconjugation followed by highly challenging purification steps and characterization. Because of the above mentioned challenges, there are no reports on semi-synthetic self-assembling protein assemblies containing an engineered disulfide bond. Although extremely challenging, it is highly important to develop a chemical method for design of this class of semi-synthetic materials because of their applicability in different fields such as diagnostics^1^ and therapeutics^2^. Considering these merits, we took up this challenge. Our ability to make monodisperse macromolecular probes containing a disulfide bond coupled with in-house developed MAPLab technology provide opportunities to design suite of redox-responsive semi-synthetic self-assembling protein assemblies.

## Results

### Macromolecular Design and Synthesis of a Redox-Sensitive Amphiphilic Activity-Based Probe

The macromolecular design has four core structural elements: (i) hydrophilic globular protein, (ii) monodisperse flexible hydrophilic linker, (iii) redox-responsive group, and (iv) monodisperse hydrophobic dendron (Figure 1).

**Figure 1.**
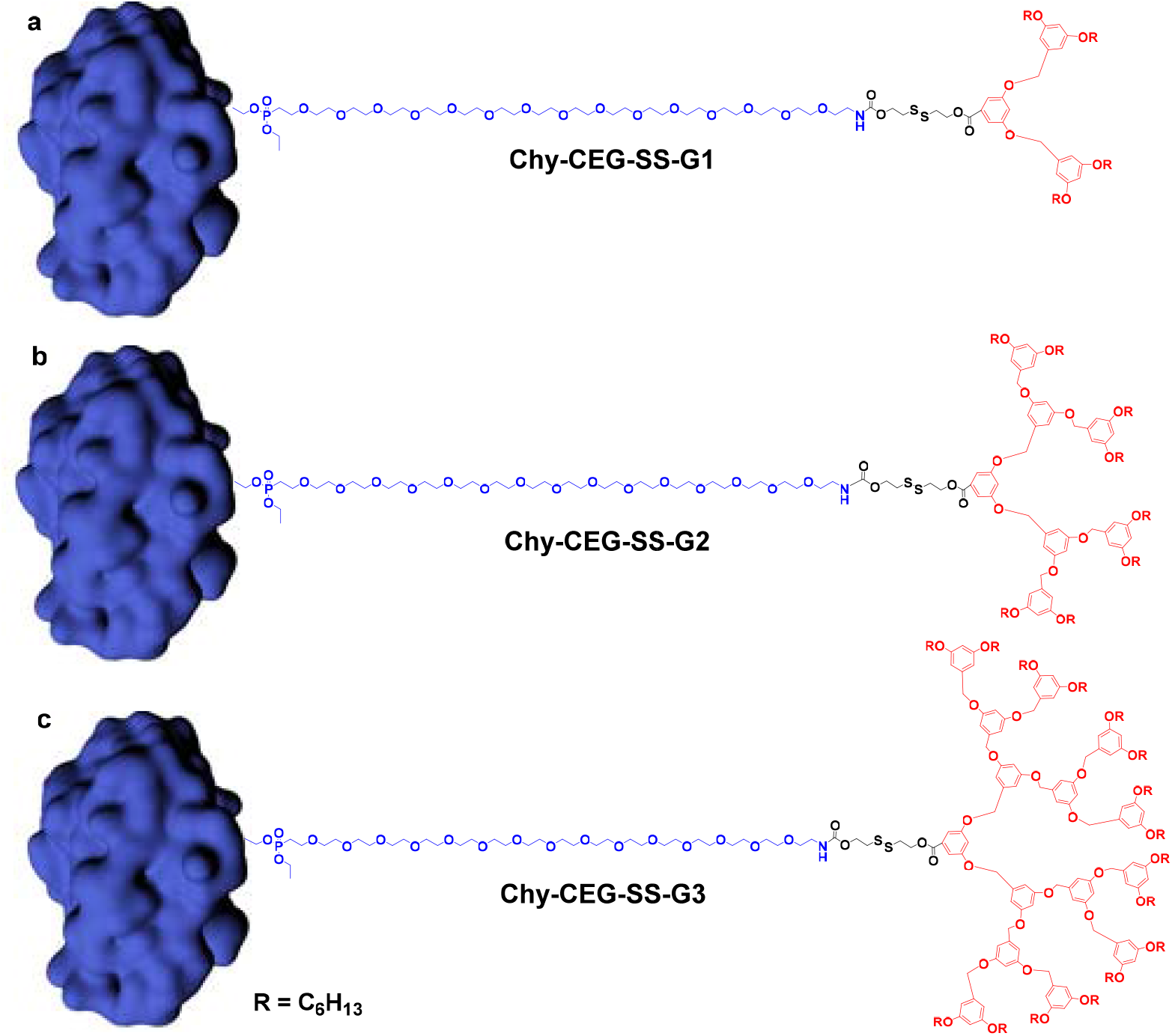
Schematic representation of chemical structures of redox-sensitive protein-dendron bioconjugates. These conjugates are composed of protein (blue = chymotrypsin), hydrophilic oligoethylene glycol linker (blue) (CEG = cetylethylene glycol), redox-sensitive disulphide bond (black), and hydrophobic dendron block (red). Structures of (a) Chy-CEG-SS-G1 (b) Chy-CEG-SS-G2 (c) Chy-CEG-SS-G3.

The design strategy for the synthesis of stimuli-responsive macromolecular probe is shown in Scheme 2a. First, we synthesized G1, G2, and G3 esters by following a previously reported method with slight modifications (Scheme 1).^14b,d^ Then, the obtained G1 (2a), G2 (3a), and G3 (4a) esters compounds were subjected to hydrolysis by treatment with NaOH in ethanol to get corresponding acid derivatives (5b). The G1-, G2-, and G3-acid derivatives (5b) were then reacted with 2,2’-disulfanediylbis(ethan-1-ol) in the presence of EDC and DMAP in DCM to get compound 5c followed by activation using N, N’-DSC, in the presence of Et_3_N to afford compound 5d. The activated ester 5d was then reacted with linker amine (6) in the presence of Et_3_N and DMF to obtain compound 5e. Then, the diphosphonate ester 5e was heated with lithium bromide (LiBr) in DMF to get monophosphonate ester 5f, which finally on fluorination using DAST in DCM afforded fluorophosphonate derivative 5g (Scheme 2).

**Scheme S1.**
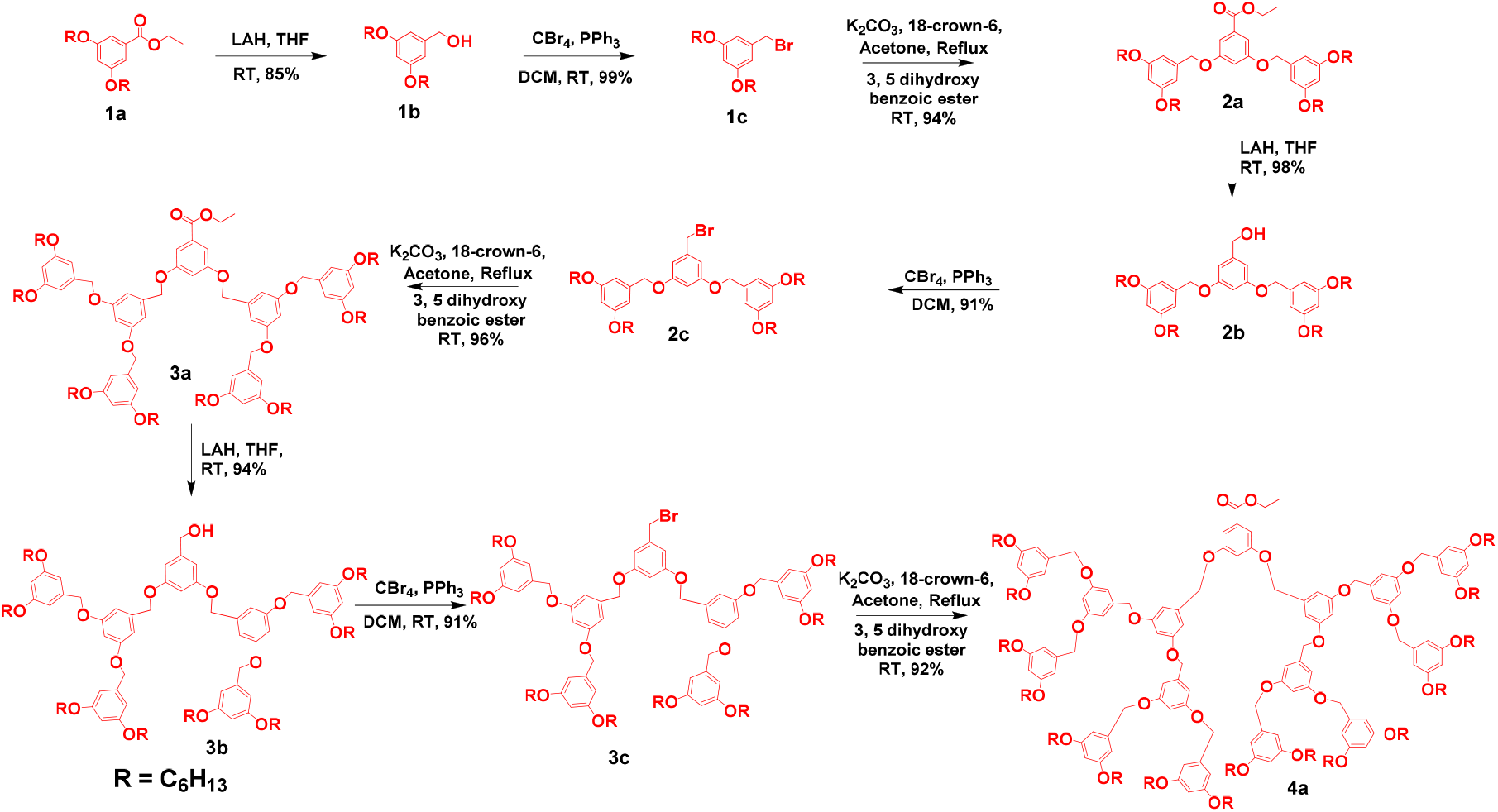
Scheme for the synthesis of ester-terminated dendrimers.

### Synthesis of Redox-Sensitive Protein-Dendron Conjugates

The conjugation of redox-sensitive macromolecular probes with chymotrypsin was attempted in 50 mM phosphate buffer pH 7.4 by using a micelle-assisted protein labelling (MAPLab) technology reported by our group. In this approach, the redox-responsive macromolecular AABPs were solubilized in 10X critical micelle concentration (CMC) of triton X-100 at room temperature. Subsequently, obtained homogeneous solution was then treated with chymotrypsin (Scheme 2b). Then, the extent of the formation of protein-dendron bioconjugates was monitored using MALDI-ToF at different time points. After 12 h, a significant peak for all the bioconjugates was observed with minor unreacted native protein (Figure 3a).

**Scheme 2.**
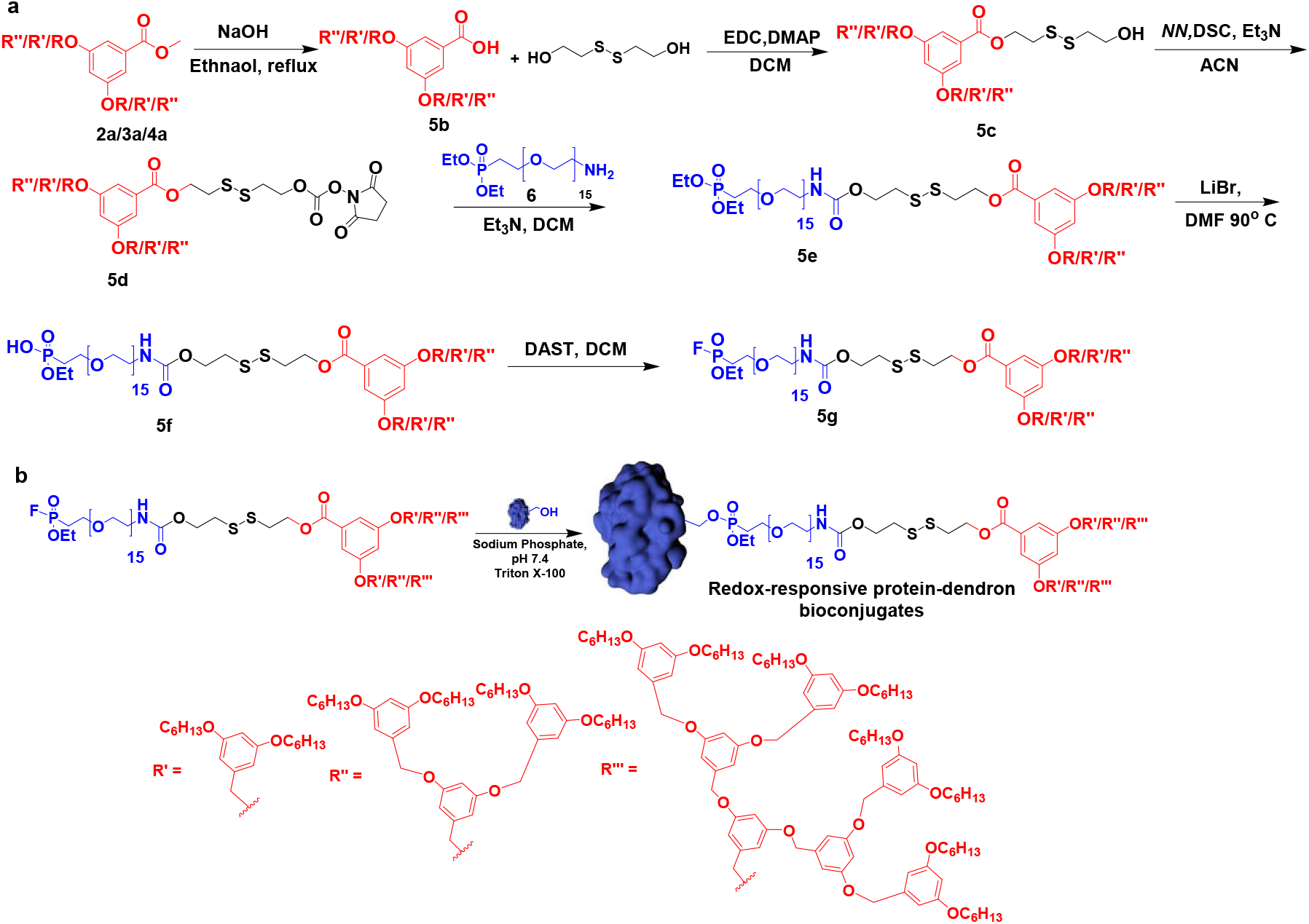
Scheme for the synthesis of redox-sensitive protein-dendron AABPs and bioconjugates. **(a)** Scheme for the synthesis of redox-sensitive macromolecular probes. **(b)** Scheme for the synthesis of redox-sensitive protein-dendron bioconjugates.

**Figure 3.**
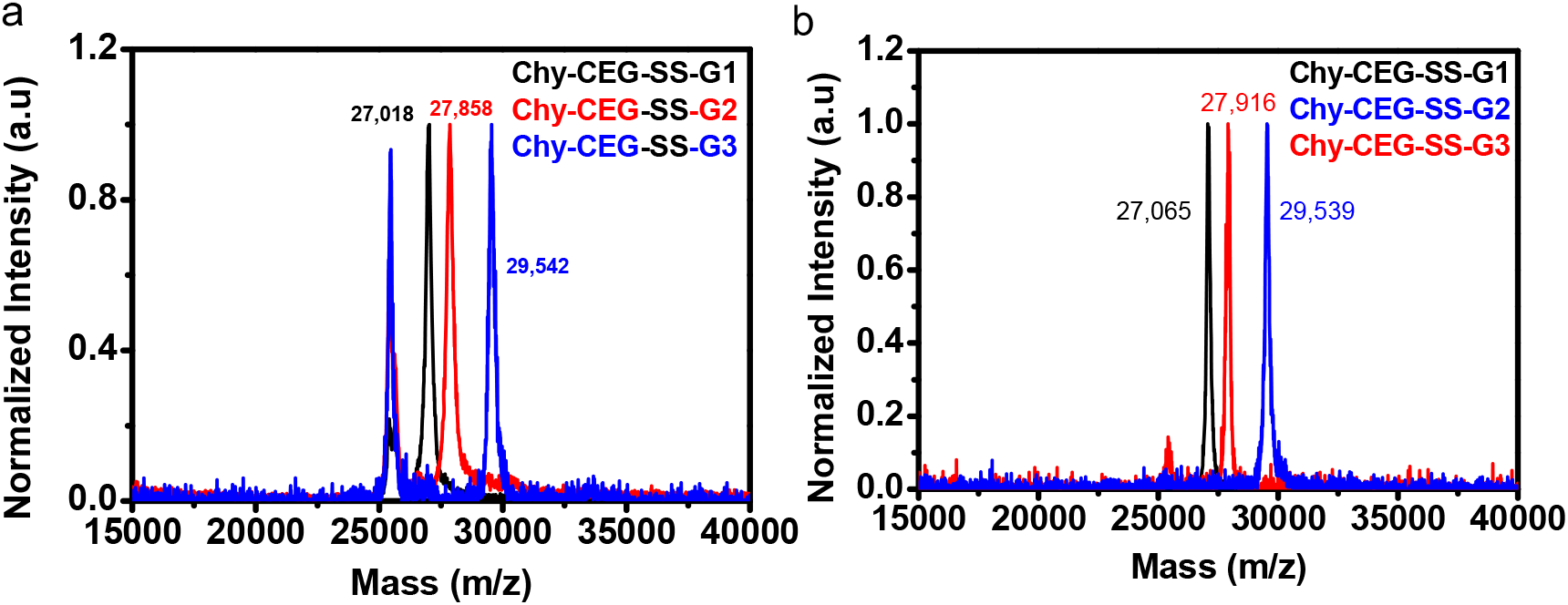
MALDI-ToF characterization redox-sensitive protein-dendron bioconjugates. **(a)** MALDI-ToF data of the reaction mixtures and **(b)** purified redox-responsive protein-dendron bioconjugates, respectively.

Next, the purification of these conjugates is performed by using the previously reported procedure by our group.^14^ Briefly, triton X-100 and excess of macromolecular probe in the reaction mixture were separated by using IEX chromatography. The unreacted protein and its corresponding conjugate were separated by using SEC chromatography, where the conjugate being facially amphiphilic forms higher-order complex driven *via* hydrophobic interaction and elutes at an earlier time point, while the monomeric native protein elutes at a later time point (Figure S1). MALDI-ToF results of purified conjugates show that they are extremely pure and devoid of any native protein (Figure 3b).

### Self-Assembly Studies of Redox-Sensitive Protein-Dendron Bioconjugate

Next, we sought to investigate the self-assembling ability of redox-sensitive protein-dendron conjugates. To do that we have carried out dynamic-light scattering and size-exclusion chromatography experiments (Figure 4a-b). All three bioconjugates exhibited monomodal distribution with very low poly-dispersity index. The hydrodynamic diameters of the Chy-CEG-SS-G1/G2/G3 complexes are found to be 12, 15, and 17 nm, respectively, indicating that the size of redox-sensitive protein-dendron complexes increases with an increase in the size of the dendron (Figure 4a). Excited by these results, we wanted to check how these bioconjugates behave in SEC. To our surprised, we observed very sharp peak for all three bioconjugates. In accordance with DLS results, Chy-CEG-SS-G3 eluted earlier (14 mL) followed by Chy-CEG-SS-G2 (15 mL) and Chy-CEG-SS-G1 (16 mL) (Figure 4b).

**Figure 4.**
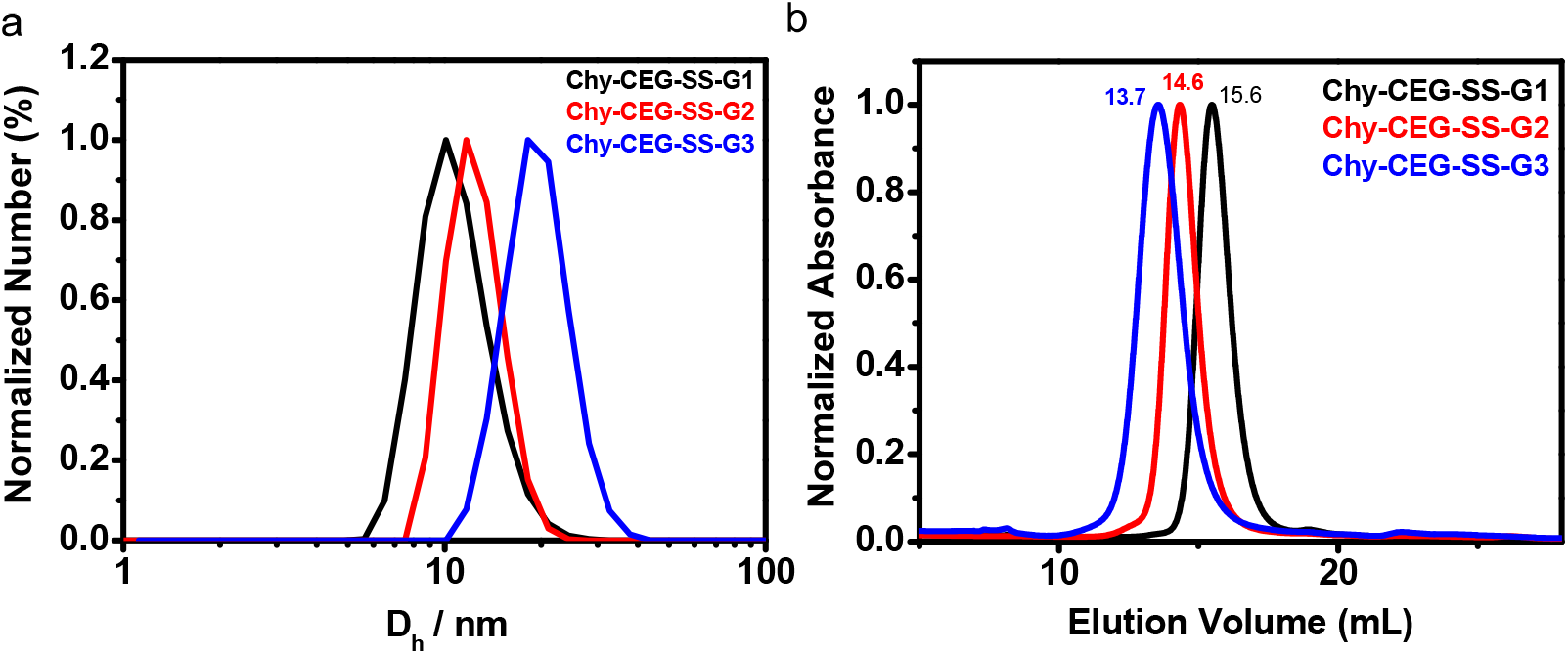
Self-assembly studies of redox-sensitive protein-dendron bioconjugates. (a) The hydrodynamic diameter of Chy-CEG-SS-G1/G2/G3 complexes is determined using DLS. (b) SEC of Chy-CEG-SS-G1/G2/G3 complexes to determine their elution volume.

Both DLS and SEC studies revealed that the incorporation of disulfide moiety does not affect the self-assembly behavior of protein dendron bioconjugates. In addition, protein complexes exhibit narrow and monomodal distribution indicative of their nearly monodisperse character. The DLS and SEC profiles of protein-dendron conjugates mostly resemble self-assembly profiles of natural and synthetic protein cages^13^ rather than protein-polymer conjugates.^12^

### Dis-Assembly Studies of Redox-Sensitive Protein-Dendron Complexes

Having established the self-assembling behavior of bioconjugates, we wanted to test whether the custom-designed protein nanoassemblies can be programmed to disassemble. Our hypothesis is upon treatment with DTT, the disulfide functionality which connects the hydrophobic dendron to the hydrophilic linker and the protein would cleave that would convert facially amphiphilic protein into hydrophilic protein, the loss of attractive hydrophobic interaction then would lead to disassembly of protein nanoassemblies (Figure 5).^14a,d^

**Figure 5.**
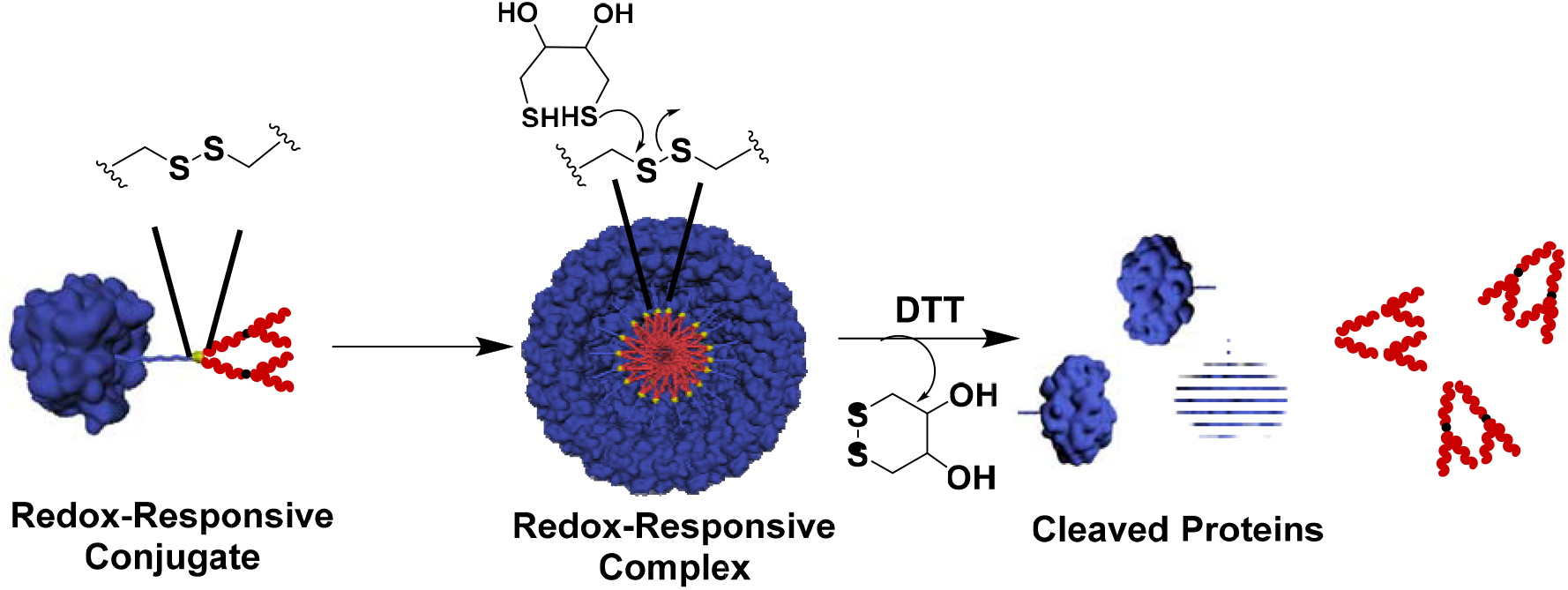
Schematic representation of self-assembly and disassembly reactions of redox-sensitive protein-dendron bioconjugates.

In order to test this hypothesis, we performed dis-assembly studies of Chy-CEG-SS-G1 in 50 mM sodium phosphate pH 7.4. The Chy-CEG-SS-G1 bioconjugate was treated with the different equivalent of DTT (10, 20, and 30 eq) for 12 hours. The DTT treated samples were then subjected to SEC in order to investigate the disassembly profile. As expected, before DTT treatment, the protein complex was intact and therefore eluted earlier (16 mL) compared to native monomeric protein (19 mL). However, the SEC of DTT incubated samples revealed that the entire complex was disassembled to its consecutive monomers with 10 eq of DTT (Figure 6a). This experiment reveals that the 10 eq. of DTT is sufficient to completely disassemble the protein complex Chy-CEG-SS-G1.

**Figure 6.**
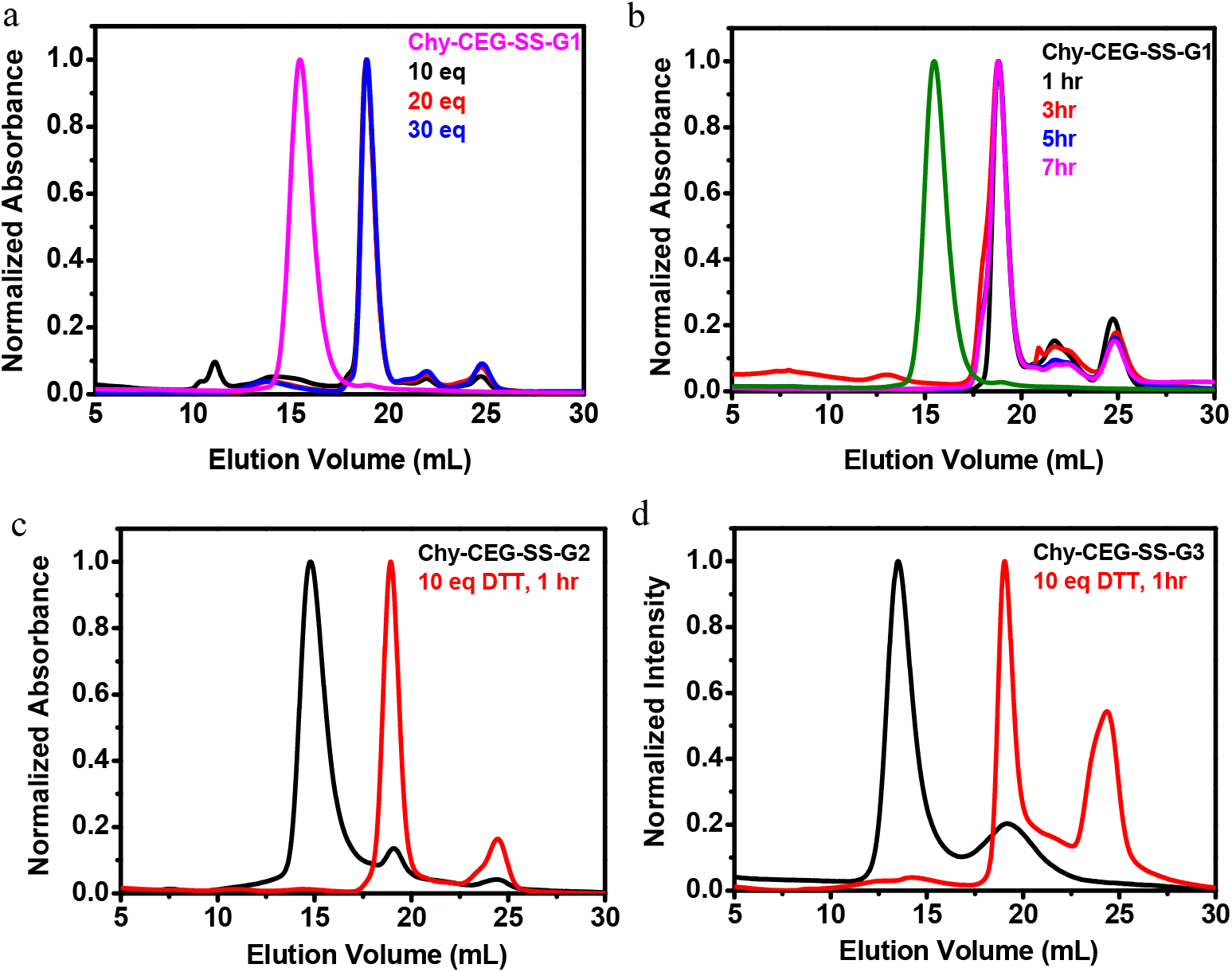
Dis-assembly profile of redox-sensitive protein-dendron protein bioconjugates; (a) Concentration dependent disassembly studies of Chy-CEG-S-S-G1, (b) Time-dependent dis-assembly studies of Chy-CEG-S-S-G1, (c) Disassembly studies of Chy-CEG-SS-G2 (d) Disassembly studies of Chy-CEG-SS-G3.

Next, in order to check whether the incubation period of DTT has any effect on disassembly behavior, the bioconjugate Chy-CEG-SS-G1 complex was incubated with 10 eq of DTT for the different incubation period. The SEC results reveal that 60 mins of incubation is sufficient to achieve the complete disassembly as evident from disappearance of peak at 16 mL (Figure 6b). After establishing the time and the concentration of DTT required to achieve the complete dis-assembly, we sought to investigate the dis-assembly behavior of the other two redox responsive protein complexes, i.e., Chy-CEG-SS-G2/G3. We were delighted to notice that both Chy-CEG-G2/G3 protein complexes also could be disassembled into respective monomeric proteins completely upon treatment with 10 eq. of DTT for about 60 mins. Based on the above experiments, it is evident that the treatment of 10 eq of DTT for 60 mins is sufficient to disassemble the Chy-CEG-SS-G1/G2G3 complexes.

## Discussion

Rational design of stimuli-responsive drug delivery systems (SRDDS) provides to opportunity for the targeted delivery of drugs to the right location.^16^ Among the various stimuli both external (temperature^17^, light^18^, sound^19^, etc) and internal (pH^20^, redox^21^, and enzyme^23^), redox-trigger is particularly attractive because of its success in clinics. Previous efforts in this area have focused on developing a chemical methodology for strategic placement of disulfide functionalities in different drug delivery systems.^4-9^ However, to our knowledge, engineering of disulfide functionality inside the interior of semi-synthetic protein assemblies have never been attempted. This is mainly attributed to the challenges associated with the system. First of all, globular proteins are hard to work with because of presence of various functional groups therefore it is important to develop a bioorthogonal reaction which can work in an aqueous medium. Secondly, the modified protein (semi-synthetic protein) should be able self-assemble into protein assemblies of defined size. Thirdly, installation of disulfide functionality should not interfere with the self-assembling ability of semi-synthetic proteins and finally designed protein assemblies should be responsive to redox trigger.

In this work, we have demonstrated the design and synthesis of monodisperse macromolecular probes containing disulfide functionality. The modular approach of our synthetic methodology allowed us to make suite of probes containing different dendrons. Further, we have used MAPLab technology^14^ for site-specific bioconjugation of chymotrypsin. The custom designed facially amphiphilic semi-synthetic proteins have found to self-assemble into protein nanoassemblies of defined size. The size of the protein nanoassemblies can be systematically tuned by choosing an appropriate dendron. The above results emphasize that the molecular design is more tolerant to stimuli-responsive functional group mutations.

Disassembly studies reveal that irrespective of protein nanoassemblies size and hydrophobic nature, all of them could be disassembled quantitatively. The above results suggest that the interiors of protein assemblies are quiet accessible as DTT can react with the engineered disulfide functionality. The result obtained from this work also reiterates our previous findings that the attractive interaction between the hydrophobic groups (benzyl-ether dendron in this case) is the major driving for the protein self-assembly.^14a,d^ Removal of hydrophobic group through the selective cleavage leads to disassembly of protein assemblies due to conversion of amphiphilic proteins into hydrophilic proteins. The result obtained in this study is quite different from our previous results obtained with photo-sensitive protein assemblies. In the previous case, we had achieved only partial disassembly because of formation of in-situ imine formation.^14d^ However, in the present case, we have observed complete disassembly because the newly formed thiol does not react with side chain of other amino present in the globular protein.

## Conclusion

In summary, we have demonstrated the successful introduction of disulfide functionality in the interiors of protein assemblies. This was achieved through modular synthesis of disulfide functionality containing macromolecular probes and successful site-specific bioconjugation utilizing a MAPLab technology. The redox-sensitive semi-synthetic proteins have shown to self-assemble into protein assemblies of defined size. The protein assemblies have programmed to disassemble upon treatment with DTT. Although the present system cannot be used for drug delivery applications because of bio-incompatibility of benzyl-ether dendron used in this study. However, it lays foundation for the design of highly defined biocompatible/biodegradable redox-responsive protein assemblies for future drug delivery applications. Those studies are under-way in our laboratories and will be reported in due course.

