## Supporting Information for "Rational Design of Programmable Monodisperse Semi-Synthetic Protein Nanomaterials Containing Engineered Disulfide Functionality"

### 1. Experimental Methods

#### 1.1. Synthesis and Characterization of Redox-Sensitive Probes and Protein-Dendron Bioconjugates.

##### 1.1.1. Protein Conjugation and Purification of Protein-Dendron Bioconjugates

The bioconjugation reaction of chymotrypsin was performed using the procedure reported by our group.[1] Briefly, 10X CMC of triton X-100 was used to solubilize redox-sensitive macromolecular AABPs (1 or 2 equivalent) and then 480 µL of 50 mM sodium phosphate pH 7.4 was added to make a final solution to 500 µL. The homogeneous solution was then treated with 500 µL of 200 µM of protein solution. This reaction mixture was then allowed to react for 24 h on rotor spin at 20 rpm at 25 ºC.

##### 1.1.2. Monitoring of Modification

Molecular weights of photo-sensitive protein-dendron conjugates were analyzed by using MALDI-ToF spectrometry. The matrix was prepared by the protocol mentioned by our group previously.[1] In brief, 2 µL of the reaction mixture or 100 µM purified protein conjugate was mixed with 2 µL of 2% TFA and 2 µL of matrix mixture, vortexed and spotted on MALDI-ToF MS plate. The plate was then loaded and fired to get accurate molecular weight both in +1 and +2 states. 100 µM protein concentration was found to be optimum for MALDI-ToF MS analysis.

##### 1.1.3. Purification of Redox-Sensitive Protein-Dendron Conjugates

All the redox-sensitive protein-dendron bioconjugates were purified by two-step purification, i.e., IEX, and SEC, performed using Akta Pure. IEX was performed to remove triton X-100 using either SP sepharose resins (GE). In cation-exchange chromatography, the column was pre-equilibrated with the same buffer (50 mM sodium phosphate pH 7.4) which was used for modification and then a sample was injected followed by post-injection equilibration for at least 2 Column Volumes (CVs) or until the complete removal of triton

X-100 for large scale reactions. The elution of native protein and its corresponding redox-sensitive protein-dendron bioconjugate together as the mixture was later achieved using 50 mM sodium phosphate pH 7.4, 1 M NaCl as elution buffer.

The obtained IEX fractions were subjected to SEC. For the separation of native protein from protein-dendron bioconjugates, 50 mM sodium phosphate pH 7.4, 200 mM NaCl was used as buffer using Superose 200 10/300, GE Healthcare column. The column was pre-equilibrated with 50 mM sodium phosphate with 200 mM NaCl for at least 2CVs, and then the sample was injected, followed by post injection equilibration with the same for at least 2CVs again or until the complete elution of the proteins.

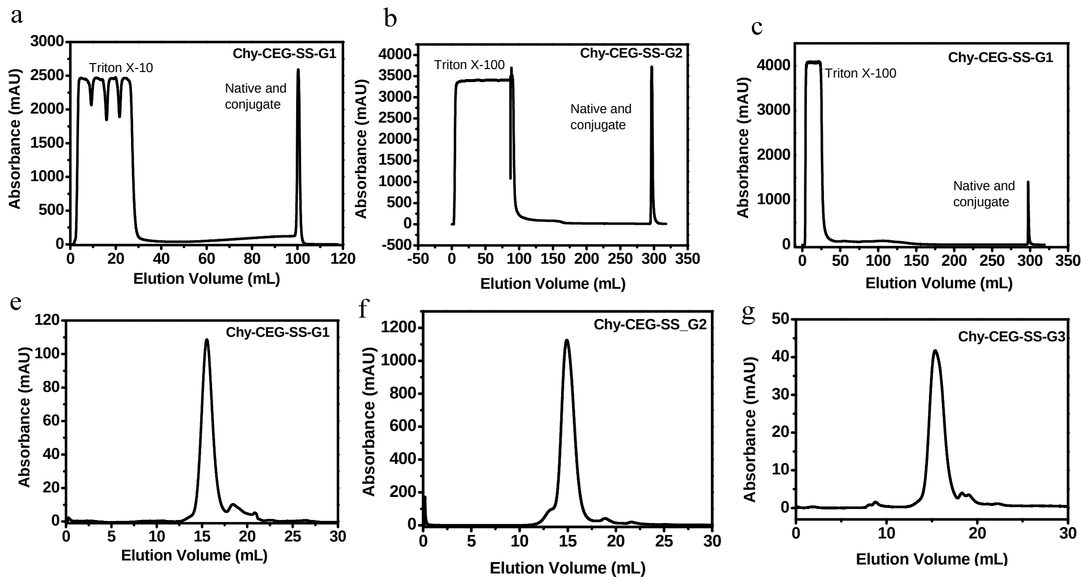


**Figure S1 | IEX and SEC chromatogram**. The IEX is performed to remove neutral triton X-100 and excess of macromolecular AABP. IEX and SEC of (a), (e) Chy-CEG-SS-G1, (b), (f) Chy-CEG-SS-G2, (c), (g) Chy-CEG-SS-G3, respectively.

#### 1.2. Self-Assembly

##### 1.2.1. Dynamic Light Scattering

The hydrodynamic diameter of redox-sensitive protein-dendron assemblies was measured using DLS (Zetasizer Nano 2590, Malvern, UK). Samples (5 mg/mL) were prepared in 50 mM sodium phosphate pH 7.4. 1 mL of sample was taken in disposable polystyrene cells, and then the mean size of the complexes was measured at 90° scattering angle.

##### 1.2.2. Size Exclusion Chromatography

The elution volume of redox-sensitive protein-dendron assemblies in SEC was determined using Akta Pure and Superose-200 10/300 GE Healthcare column. In order to do that, IEX fractions of bioconjugates were first subjected to the SEC in order to separate native protein from bioconjugates. Then, the 500 µL of the center fraction of the complex peak was again subjected to SEC by using the pre-equilibrated column mentioned above in 50 mM sodium phosphate pH 7.4, 200 mM NaCl.

#### 1.3. Programmed Dis-Assembly of a Redox-Sensitive Supramolecular Protein Complex

The redox-responsive protein complex dissolved in 50 mM sodium phosphate pH 7.4 was treated with the different equivalent of aq. DTT for 12 hours. Then, 500 µL of this was then subjected to SEC by using Superose column (pre-equilibrated sodium phosphate pH 7.4, 200 mM NaCl) and sodium phosphate pH 7.4, 200 mM NaCl as an elution buffer. By using the same protocol, the complex was incubated with 10 eq of DTT for different time points. Then the sample was subjected to the SEC using the same column and buffer mentioned above.

#### 1.4. Synthesis and Characterization of Intermediates and Redox-Sensitive probes

##### 1.4.1. General

All reagents were obtained commercially unless and otherwise stated. Reactions were performed in an oven-dried round bottom flask (RBF) and under nitrogen atmosphere. Air and moisture sensitive solvents were transferred *via* syringe. Reactions were monitored by thin-layer chromatography (TLC), and the developed chromatogram was visualized by ultraviolet (UV) lamp or by phosphomolybdic acid (PMA) staining. Product purification was accomplished by 100-200 mesh size silica gel column chromatography.

All the compounds were characterized by $^{1}H$, $^{13}C$, and $^{19}F$ (in case of fluorinated compounds) nuclear magnetic resonance (NMR) using Bruker or Jeol 400 MHz. $^{1}H$ and $^{19}F$ were recorded at an operating frequency of 400 MHz and 100 MHz for $^{13}C$ using, using TMS as an internal standard. All the $^{13}C$ Chemical shifts were mentioned in parts per million (PPM) and measured relative to residual $CHCl_3$, $CH_3OH$, or $CH_3CN$ in their deuterated solvent. Coupling constants were reported in Hertz (Hz). Multiplicities were explained as s = singlet, d = doublet, t = triplet, q = quartet, m = multiplet, quint = quintet. Mass spectra were obtained with either the MALDI-TOF MS or HRMS. Room temperature varied between 21-35 °C.

#### 1.4.3. Synthesis of monodisperse hydrophobic dendrimers

The synthesis of monodisperse hydrophilic dendrimer (2a, 3a, 4a) (Figure S1) is carried out by procedure reported by our group.[2]

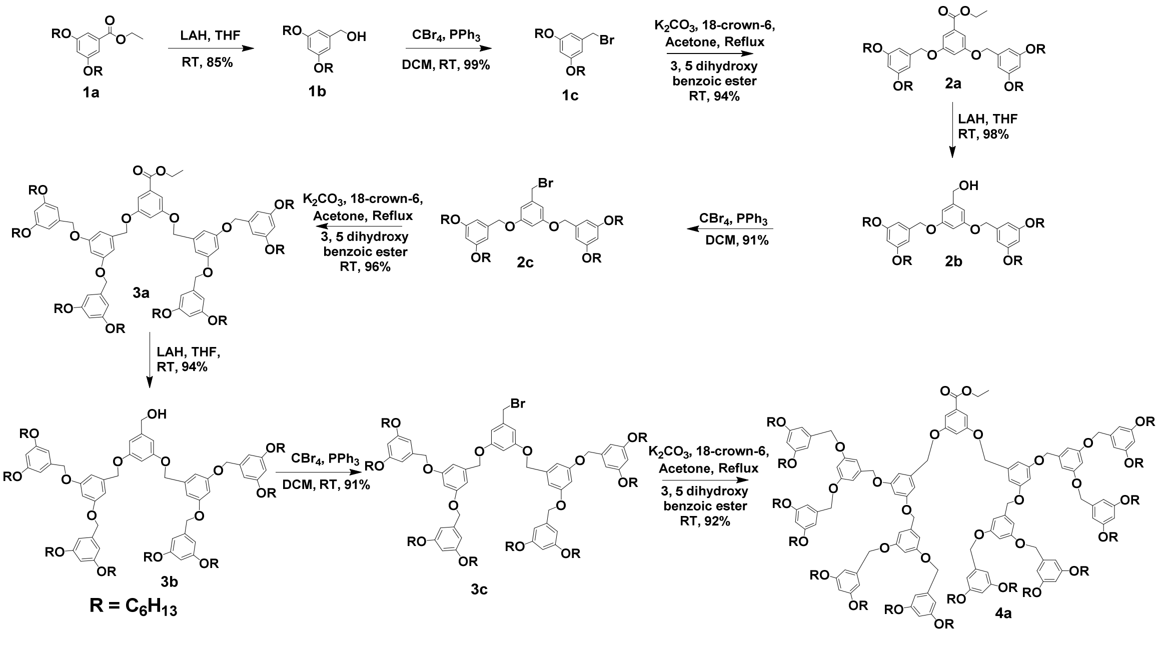


**Scheme S1 |** Scheme for the synthesis of ester-terminated dendrimers.

#### 1.4.2. Synthesis of monodisperse hydrophilic linker

The synthesis of monodisperse hydrophilic linker (6) is carried out by procedure reported by our group.[3]

##### 1.4.4. Synthesis Redox-Sensitive probes

##### General Procedure for the Synthesis of Acid - Procedure A

The ester (1 eq), sodium hydroxide (NaOH) (4 eq) were dissolved in ethanol. The mixture was then refluxed for 4 hours. Upon completion, the reaction was quenched with dropwise addition of water and acidified with conc. HCl. The obtained precipitate was filtered and washed with ethanol for two times. The combined organic layer dried over $Na_2SO_4$ and concentrated under reduced pressure to get crude, which was utilized for the next reaction without further purification.

##### General Procedure for the Synthesis of Alcohol- Procedure B

In an oven-dried RBF above-obtained acid (1 eq), 2,2'-disulfanediylbis(ethan-1-ol) (2 eq) and DMAP (0.5 eq) were taken and dissolved in DCM under stirring. Then, the solution of EDC in DCM (2 eq) was added slowly to the above mixture and allowed to stir for 12 hours. Upon completion, the reaction was quenched with water, and the resulting content was extracted in DCM thrice. Combined organic layers was dried over $Na_2SO_4$ and concentrated under reduced pressure to get the crude product, which was purified using silica gel column chromatography.

##### General Procedure for the Synthesis of Activated Ester- Procedure C

In an oven-dried RBF, above-obtained alcohol (1 eq) and N, N′-DSC (5 eq) were dissolved in ACN under stirring. Then, $Et_3N$ (5 eq) was then added slowly at RT and allowed to react for 12 h. Upon completion of the reaction, ACN and $Et_3N$ were evaporated under vacuum. The obtained residue was directly purified using silica gel column chromatography using ethyl acetate/hexane as eluent.

##### General Procedure for the Synthesis of Diphosphonate Ester - Procedure D

In an oven-dried RBF, the above obtained activates ester (1.1 eq) and amine-terminated cetylethylene glycol (1 eq) (6) were dissolved in DMF under stirring. Then, $Et_3N$ (1.1 eq) was added slowly to the reaction mixture and stirred at RT for 12 h. Upon completion of the reaction, DMF and $Et_3N$ were evaporated under vacuum. To the obtained residue, water was added and extracted thrice with DCM. The combined organic layer was dried over $Na_2SO_4$ and concentrated under vacuum to get the crude product, which was purified using silica gel column chromatography using MeOH / DCM as eluent.

#### General Procedure for the Synthesis of Monophosphonate Ester - Procedure E

In an oven-dried RBF, the above-obtained diphosphonate ester (1 eq) was taken, and LiBr (20 eq) was added. To the mixture, DMF was added and heated at 95 °C for 20 h. Upon completion, water was added and extracted thrice with ethyl acetate. The water layer was collected, and 2N HCl was added and stirred for another 30 minutes. The mixture was then extracted thrice with ethyl acetate. The combined organic layer was dried over $Na_2SO_4$ and concentrated under vacuum to get the crude product, which was used without purification.

#### General Procedure for the Synthesis of Flurophosphonate - Procedure F

To the stirring solution of the above obtained monophosphonate ester (1 eq) in DCM, DAST (4 eq) was added dropwise at -78 °C and allowed to react for 15 minutes. Upon completion of the reaction, excess of DAST and DCM were evaporated under vacuum. To the obtained residue, water was added and stirred for 2 more minutes to quench any residual DAST. The reaction mixture was then extracted thrice with DCM. The combined organic layer was dried over $Na_2SO_4$ and concentrated under vacuum to get the crude product, which was used without purification.

#### Synthesis of Photo-Sensitive Macromolecular AABP and Their Intermediates

##### Synthesis of compound 6a

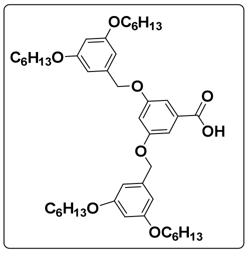

The compound **6a** was prepared by general procedure A, starting from compound 2a (3 g, 4 mmol), NaOH (0.6 g, 16 mmol). The product obtained as pale solid (2.5 g, 3.5 mmol, 87%), $R_f$=0.52 in 20% ethyl acetate / hexane. **$^1$H NMR** (400MHz, $CDCl_3$): $\delta_H$ 7.34 (d, *J*=8.9, 2.8 Hz, 1H), 6.56-6.40 (m, 8H), 5.12 (s, 2H), 4.99 (s, 4H), 3.96 (t, *J*=6.4Hz, 4H), 1.77-1.53 (m, 8H), 1.53-1.26 (m, 28H), 0.90 (t, *J*=7.2Hz, 12H). **$^{13}$C NMR (**100MHz, $CDCl_3$): 166.44, 160.67, 159.87, 138.74, 132.51, 108.52, 107.10, 105.90, 101.05, 70.46, 68.22, 61.28, 31.72, 29.36, 25.87, 22.75, 14.44, 14.26.

**Synthesis of compound 6b**

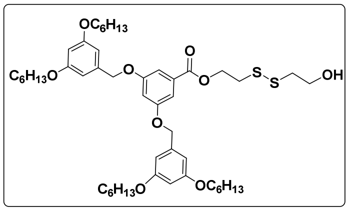

The compound **6b** was prepared by general procedure B, starting from compound **6a** (2 g, 2 mmol), 2,2'-disulfanediylbis(ethan-1-ol) ( 0.6 g, 4 mmol). The product obtained as a pale yellow solid (0.9 g, 1 mmol, 50%), $R_f$=0.20 in 20% ethyl acetate / hexane.

**Synthesis of compound 6c**

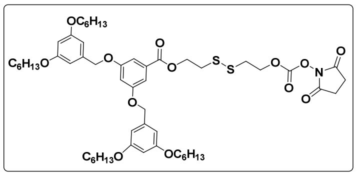

The compound **6c** was prepared by general procedure C, starting from compound **6b** (1.5 g, 2 mmol) and *N,N'*-DSC (2.5 g, 10 mmol) and, $Et_3N$ (1 g, 10 mmol). The product obtained as a pale yellow solid (1 g, 1 mmol, 60%), $R_f$=0.18 in 25% ethyl acetate / hexane.

**Synthesis of compound 6d**

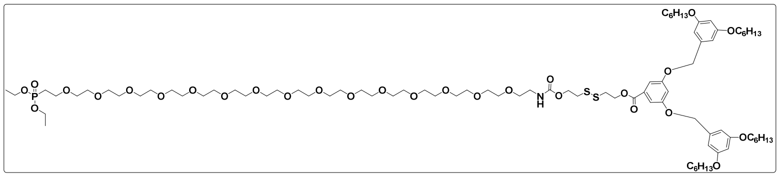

The compound **6d** was prepared by general procedure D, starting from compound **6c** (0.7 g, 0.7 mmol), compound **6** (Synthesis and characterization data for compound 6 is mentioned in chapter 3) (0.6 g, 0.7 mmol) and, $Et_3N$ (0.06 g, 0.6 mmol). The obtained product was pale yellow liquid (0.78 g, 0.4 mmol, 65%), $R_f$=0.40 in 5% MeOH / DCM. **$^1$H NMR** (400MHz, $CDCl_3$) 7.10 (dd, *J*=8.9, 2.8 Hz, 1H), 6.5. (d, m, 9H), 5.46 (s, 2H), 5.23 (s, 2H), 4.24 (m, 4H), 4.07 (t, *J*=6.4Hz, 8H), 3.77-3.52 (m, 57H), 2.19-2.05 (m, 2H), 1.85-1.60 (m, 12H), 1.44-1.29 (m, 48H), 0.82 (t, J=6.8Hz, 9H).

**Synthesis of compound 6e**

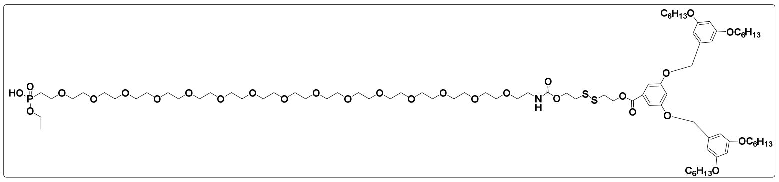

The compound **6e** was prepared by general procedure E, starting from compound **6d** (0.5 g, 0.3 mmol) and, LiBr (0.3 g, 3 mmol) was added. The obtained product was carried to the next step without further purification.

**Synthesis of compound 6f**

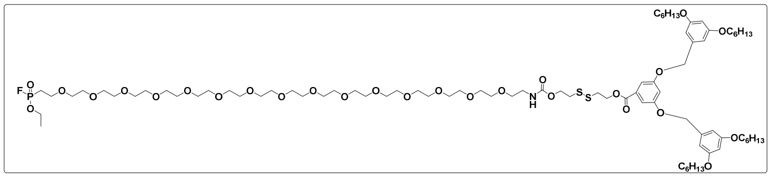

The compound **6f** was prepared by general procedure F, starting from compound **6e** (0.1 g, 0.050 mmol), DAST (0.040 g, 0.2 mmol). The product was utilized for conjugation without further purification. **$^{19}$F NMR** (400MHz, $CDCl_3$): $\delta_F$ -59.91, -62.74.

**Synthesis of compound 7a**

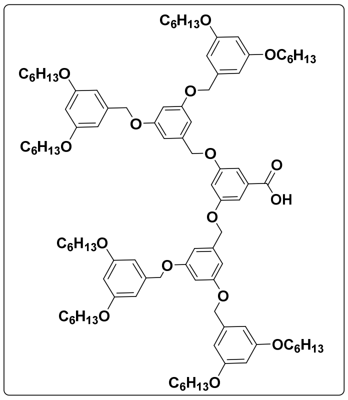


The compound **7a** was prepared by general procedure A, starting from compound **4a** (0.018 g, 0.10 mmol), NaOH (0.17 g, 0.10 mmol). The product obtained as pale solid (0.10 g, 0.06 mmol, 65%), $R_f$=0.52 in 20% ethyl acetate / hexane. **$^1$H NMR** (400MHz, $CDCl_3$): $\delta_H$ 7.33 (d, *J*=8.9, 2H), 6.71-6.43 (m, 28H), 5.00-4.94 (m, 20H), 3.97 (t, *J*=6.4Hz, 26H), 1.80-1.75 (m, 28H), 1.54-1.30 (m, 90H), 0.93 (t, *J*=7.2Hz, 44H).

**Synthesis of compound 7b**

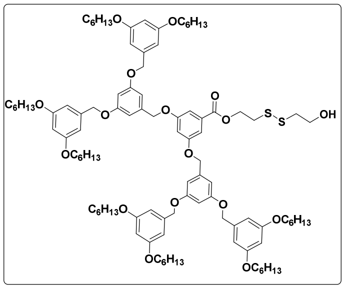

The compound **7b** was prepared by general procedure B, starting from compound **7a** (1 g, 0.6 mmol), 2,2'-disulfanediylbis(ethan-1-ol) (0.2 g, 1.2 mmol). The product obtained as a pale yellow solid (0.4 g, 0.2 mmol, 35%), $R_f$=0.20 in 20% ethyl acetate / hexane. **$^1$H NMR** (400MHz, $CDCl_3$): $\delta_H$ 7.29 (d, *J* = 2.8 Hz, 2H), 6.62-6.38 (m, 21H), 5.61 (s, 2H), 5.03-4.89 (m, 12H), 3.94 (t, *J*=6.4Hz, 14H), 3.06-2.91 (m, 4H), 1.85-1.69 (m, 16H), 1.47-1.22 (m, 53H), 0.90 (t, *J*=7.2Hz, 24H).

**Synthesis of compound 7c**

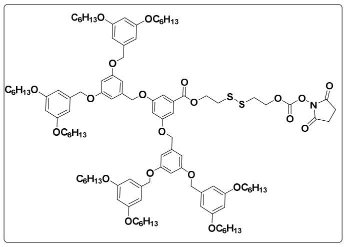

The compound **7c** was prepared by general procedure C, starting from compound **7b** (0.4 g, 0.2 mmol), *N,N′*-DSC (0.3 g, 1 mmol) and $Et_3N$ (0.1 g, 1 mmol). The product obtained as a pale yellow solid (0.3 g, 0.1 mmol, 66%), $R_f$=0.18 in 25% ethyl acetate / hexane.

##### Synthesis of compound 7d

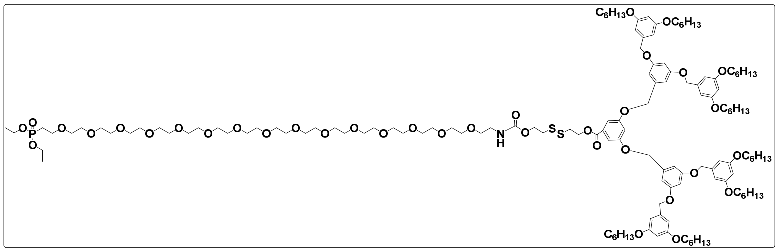

The compound **7d** was prepared by general procedure D, starting from compound **7c** (0.3 g, 0.1 mmol), compound **6** (Synthesis and characterization data for compound 6 is mentioned in chapter 3) (0.09 g, 0.1 mmol) and, $Et_3N$ (0.01 g, 1 mmol). The product obtained was pale yellow liquid (0.3 g, 0.1 mmol, 70%), $R_f$=0.40 in 5% MeOH / DCM. **$^1$H NMR** (400MHz, $CDCl_3$): $\delta_H$ 7.17 (d, 2.8 Hz, 1H), 6.64-6.40 (m, 21H), 5.0-4.82 (m, 12H), 4.20-410 (m, 20H), 3.70-3.45 (m, 66H), 2.19-2.06 (m, 2H), 1.86-1.66 (m, 18H), 1.50-1.18 (m, 70H), 0.89 (t, *J*=7.2Hz, 30H). **$^{13}$C NMR (**100MHz, $CDCl_3$):

##### Synthesis of compound 7e

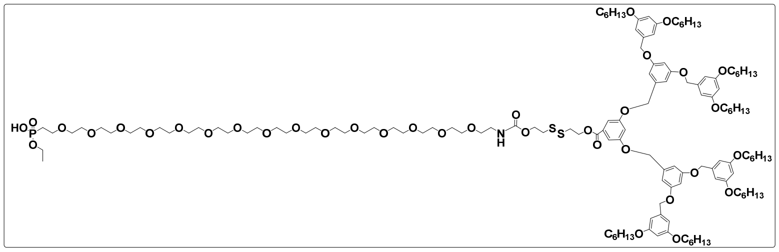

The compound **7e** was prepared by general procedure E, starting from compound **7d** (0.140 g, 0.02 mmol) and, LiBr (0.080 g, 0.5 mmol). The obtained product was carried to the next step without further purification.

#### Synthesis of compound 7f

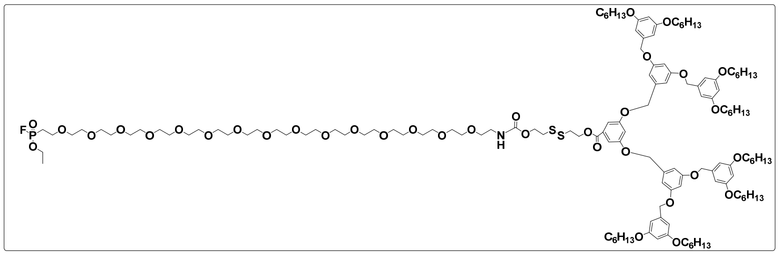

The compound **7f** was prepared by general procedure F, starting from compound **7e** (0.08 g, 0.01 mmol), DAST (0.02 g, 0.05 mmol). The obtained product was utilized for conjugation without further purification. **$^{19}$F NMR** (400MHz, $CDCl_3$): $\delta_F$ -59.91, -62.74.

#### Synthesis of compound 8a

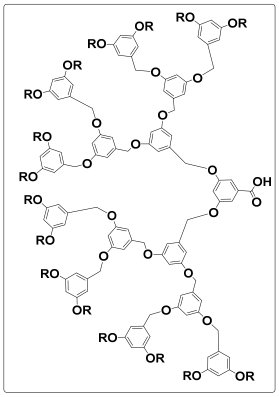

The compound **8a** was prepared by general procedure A, starting from compound **6a** (3 g, 0.9 mmol), and NaOH (0.15 g, 3.6 mmol). The product obtained as pale solid was carried to the next step without purification. **$^1$H NMR** (400MHz, $CDCl_3$): $\delta_H$ 7.42 (d, $J$ = 2.4 Hz, 2H), 6.73-6.45 (m, 45H), 5.05-4.90 (m, 28H), 3.96 (t, $J$ = 6.8Hz, 34H), 1.83-1.71 (m, 34), 1.49-1.22 (m, 134H), 0.94 (t, $J$ = 7.2Hz, 54H).

#### Synthesis of compound 8b

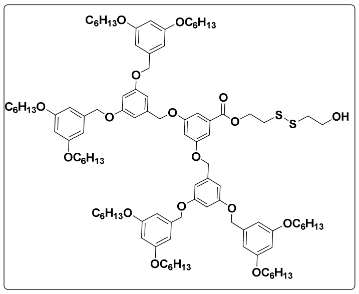

The compound **8b** was prepared by general procedure B, starting from compound **8a** (2 g, 0.6 mmol), 2,2'-disulfanediylbis(ethan-1-ol) (0.2 mg, 1.2 mmol). The product obtained as a pale yellow solid (0.7 g, 0.2 mmol, 36%), $R_f$=0.20 in 20% ethyl acetate / hexane. **$^1$H NMR** (400MHz, $CDCl_3$): $\delta_H$ 7.30 (d, $J$ = 2.4Hz, 2H), 6.69-6.37 (m, 45H), 5.01-4.94 (m, 28H), 3.92 (t, $J$ = 6.8Hz, 34H), 3.04-2.85 (m, 4H), 1.82-0.86 (m, 270H).

**Synthesis of compound 8c**

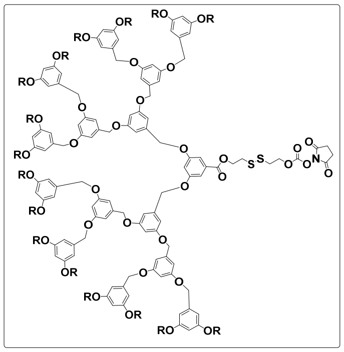

To the mixture of compound **7b** (0.7 g, 0.2 mmol) and pyridine (0.1 g, 1 mmol) in DCM, a solution of 4-nitrophenyl chloroformate (0.2 g, 1 mmol) in DCM was added at 0 °C. The mixture was allowed to react for 3 hours at RT. Then, the solvent was evaporated under reduced pressure to get the crude product, which was then purified using silica gel column chromatography (0.35 g, 0.1 mmol, 55%), $R_f$=0.20 in 20% ethyl acetate/hexane.

**Synthesis of compound 8d**

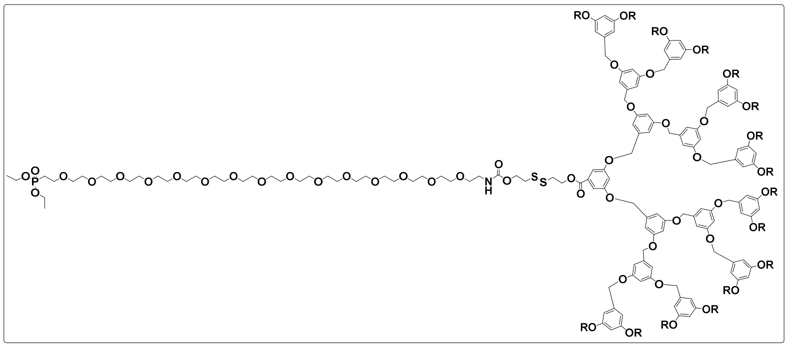

The compound 8**d** was prepared by general procedure D, starting from compound **8c** (0.4 g, 0.1 mmol), compound **6** (Synthesis and characterization data for compound 6 is mentioned in chapter 3) (0.09 g, 0.1 mmol) and, $Et_3N$ (0.03 g, 0.3 mmol). The product obtained was pale yellow liquid (0.2 g, 0.07 mmol, 65%), **$^1$H NMR** (400MHz, $CDCl_3$): $\delta_H$

7.3 (d, $J$ = 2.4Hz, 2H), 6.66-6.38 (m, 45H), 5.30 (3, 14H), 4.95-4.93 (m, 26H), 4.20-4.15 (m, 4H), 4.02-3.93 (m, 30H), 3.91-3.76 (m, 70H), 2.19-2.09 (m, 2H), 1.78-1.71 (m, 34H), 1.51-1.26 (m, 144H), 0.89 (t, $J$=6.4Hz, 58H).

**Synthesis of compound 8e**

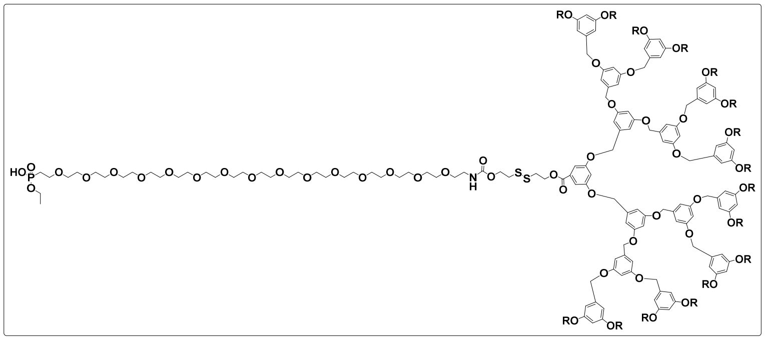

The compound **8e** was prepared by general procedure E, starting from compound **8d** (0.6 g, 0.04 mmol) and LiBr (0.12 g, 0.5 mmol). The obtained product was carried to the next step without further purification.

**Synthesis of compound 8f**

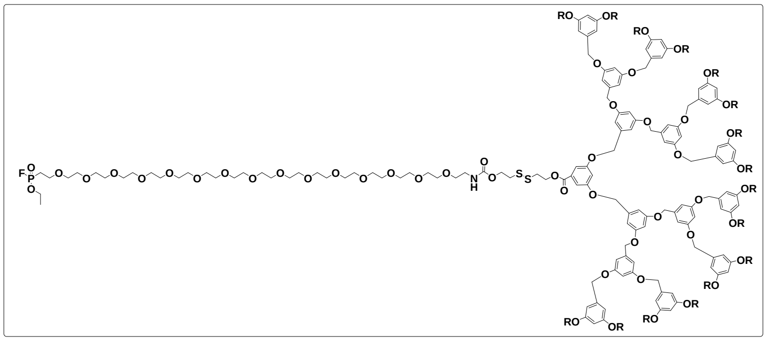

The compound **8f** was prepared by general procedure H, starting from compound **8e** (0.1 g, 0.04 mmol), DAST (0.005 g, 0.05 mmol). The obtained product was utilized for conjugation without further purification. $^{19}$**F NMR** (400MHz, $CDCl_3$): $\delta_F$ -59.91, -62.74.

#### 2. Appendix III Characterization data of synthesized compounds

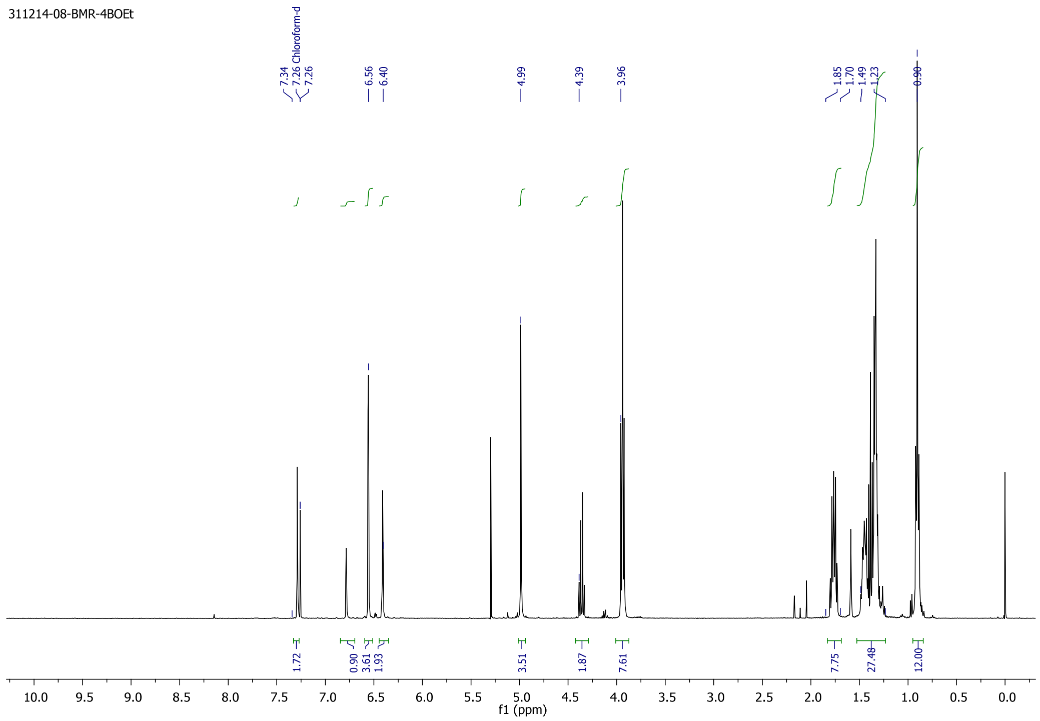


**$^1$H NMR of compound 6a**

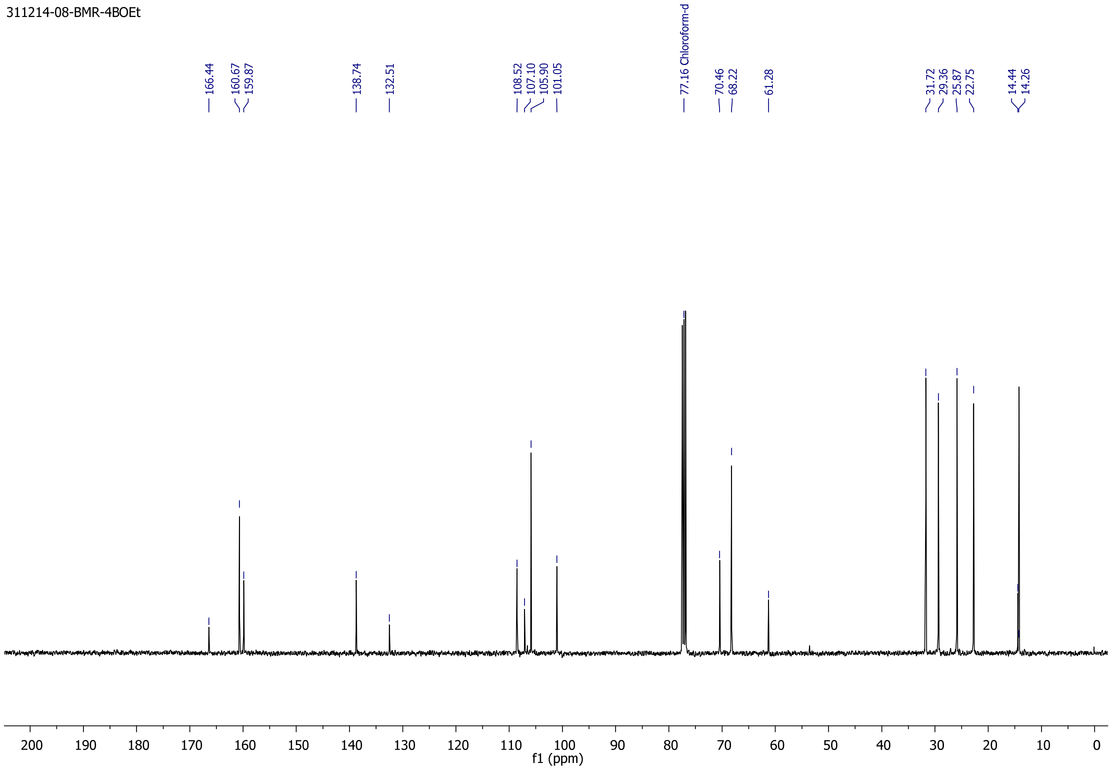


**$^{13}$C NMR of compound 6a**

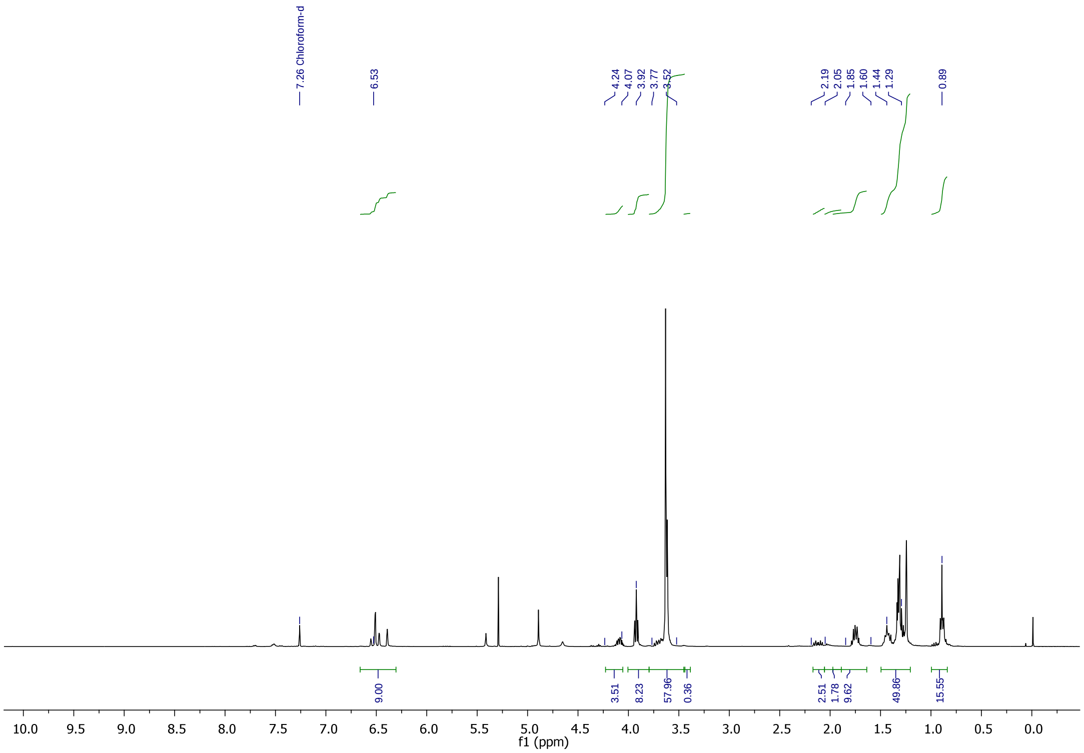


### [1]H NMR spectrum of compound 6d

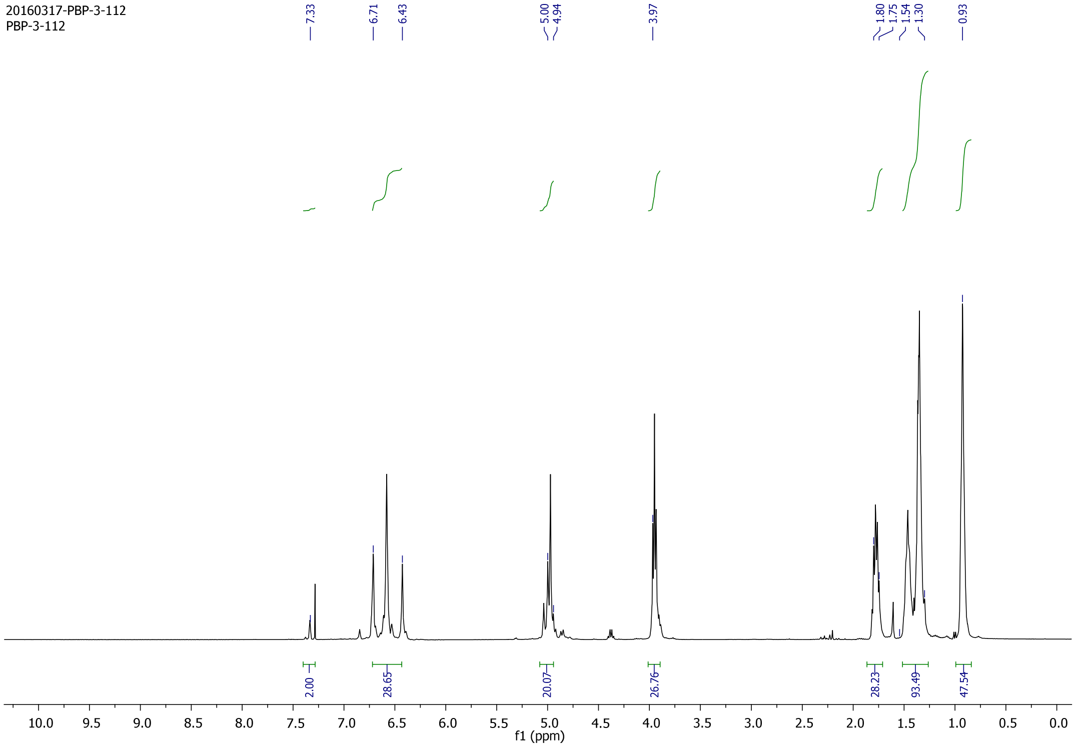


### [1]H NMR of compound 7a

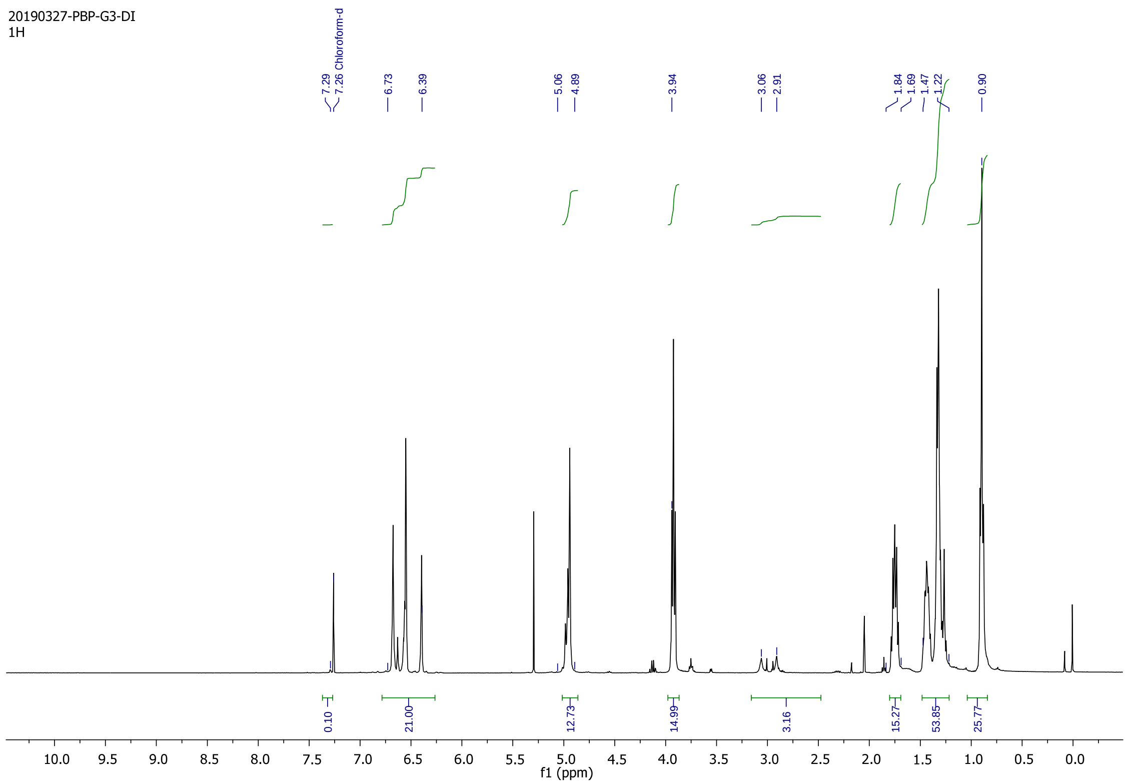


### [1]H NMR of compound 7b

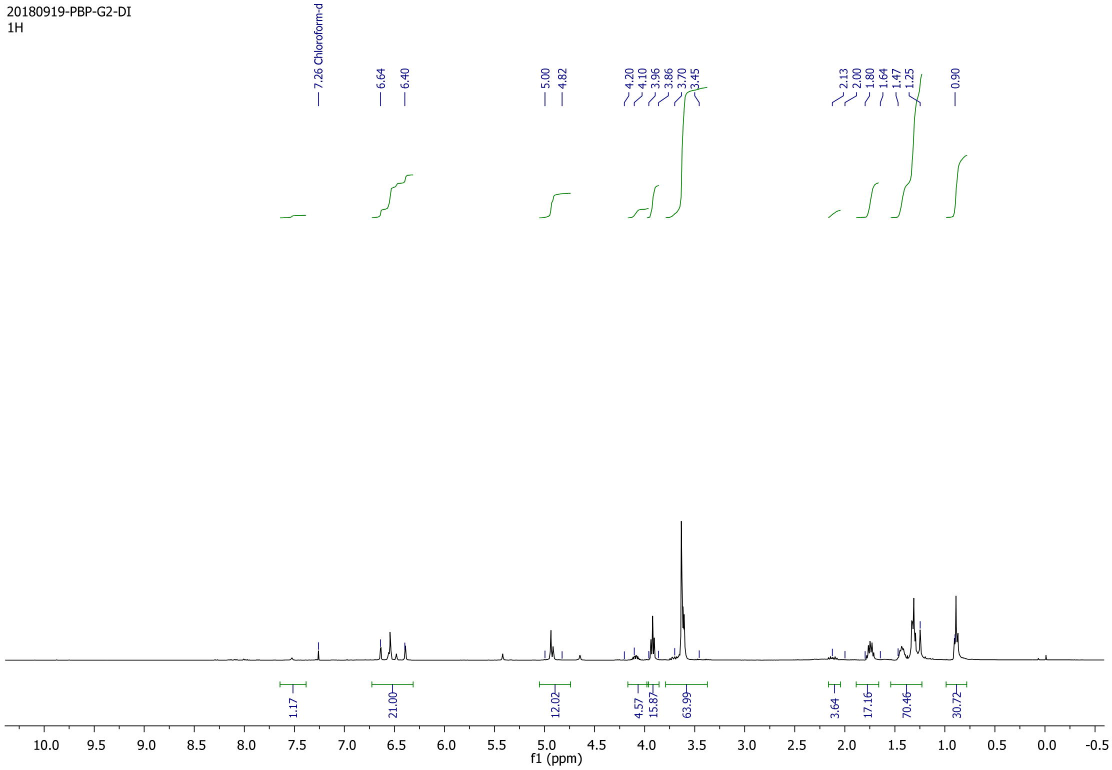


### [1]H NMR of compound 7d

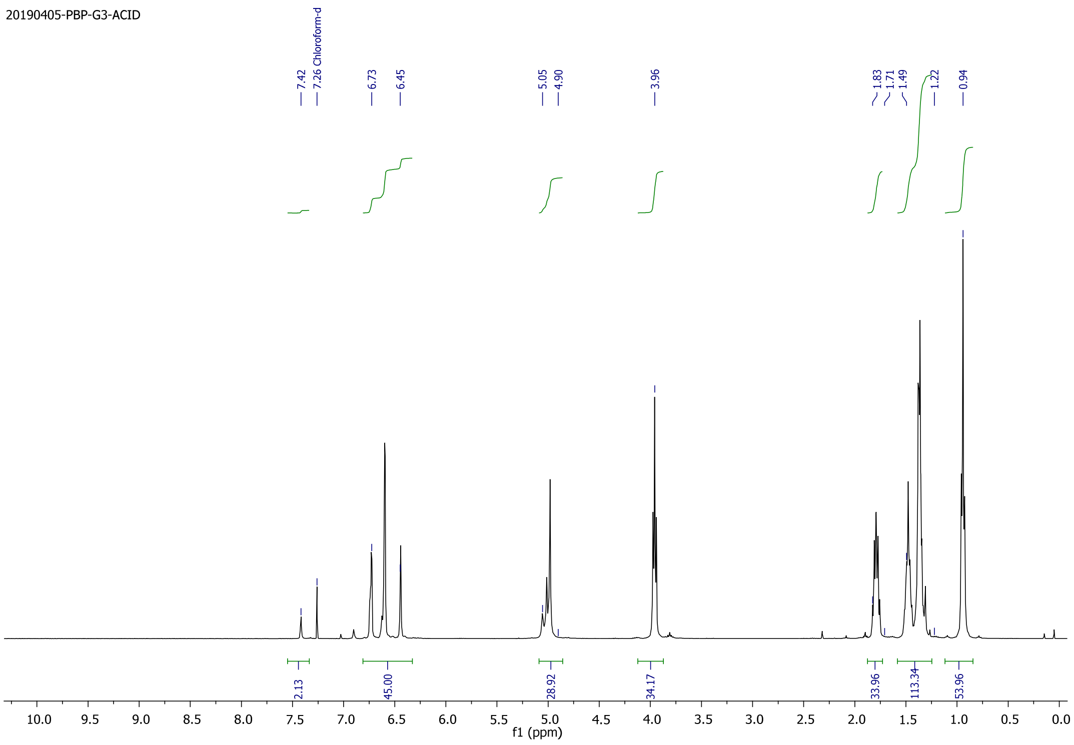


### [1]H NMR of compound 8a

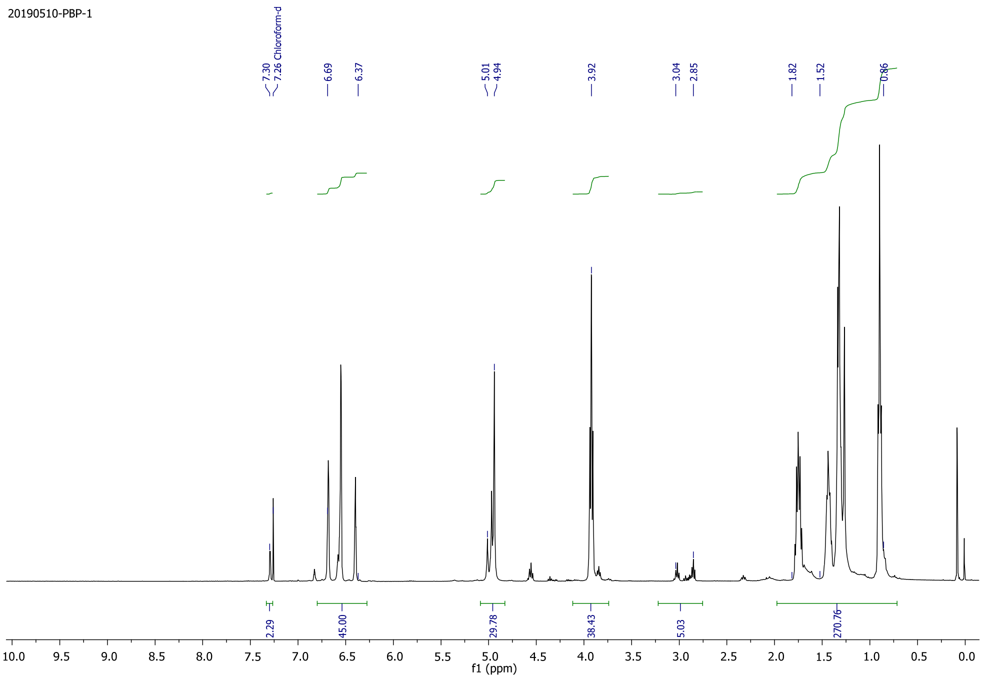


### [1]H NMR of compound 8b

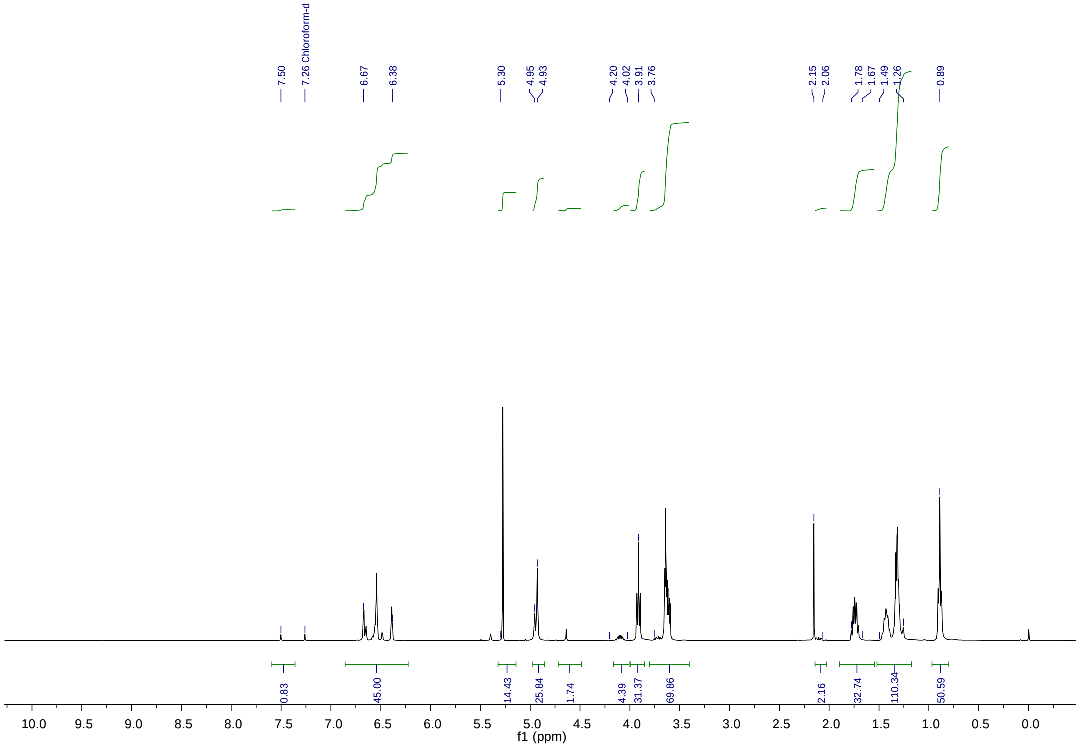


**$^{1}$H NMR of compound 8d**
